# ORIGINS: a protein network-based approach to quantify cell pluripotency from scRNA-seq data

**DOI:** 10.1101/2022.05.09.491232

**Authors:** Daniela Senra, Nara Guisoni, Luis Diambra

## Abstract

Trajectory inference is a common application of scRNA-seq data. However, it is often necessary to previously determine the origin of the trajectories, the stem or progenitor cells. In this work, we propose a computational tool to quantify pluripotency from single cell transcriptomics data. This approach uses the protein-protein interaction (PPI) network associated with the differentiation process as a scaffold and the gene expression matrix to calculate a score that we call differentiation activity. This score reflects how active the differentiation network is for each cell. We benchmark the performance of our algorithm with two previously published tools, LandSCENT [1] and CytoTRACE [2], for four data sets: breast, colon, hematopoietic and lung. We show that our algorithm is more efficient than LandSCENT and requires less RAM memory than the other programs. We also illustrate a complete workflow from the count matrix to trajectory inference using the breast data set.

**Graphical abstract:** 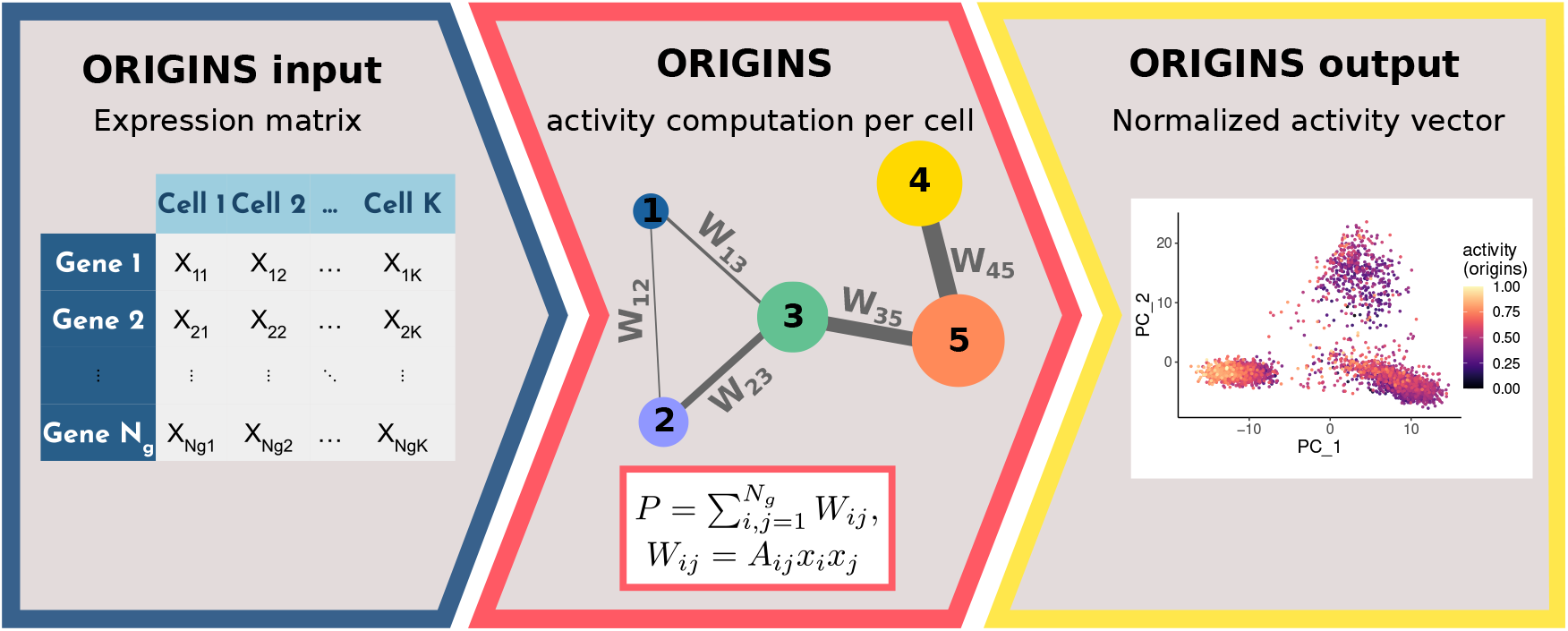

## Method details

### Background

Recent advances in single cell RNA sequencing (scRNA-seq), that allow the transcriptional profiling of single cells, offer a promising capability to explain developmental processes. The ability to quantify pluripotency is relevant to comprehend differentiation, cell lineages and lineage hierarchy. This task may also be important for cancer research to identify cancer stem cells, which have been suggested as responsible for metastasis, remission and resistance to therapies [3]. It may also be a critical step to perform trajectory inference, a popular application of single cell transcriptomics to unveil differentiation processes.

Several algorithms were developed to reconstruct differentiation pathways by using scRNA-seq. In 2019 Saelens *et al*. reported the existence of more than 70 trajectory analysis techniques [4] and in the recent years many more have emerged [5, 6, 7]. Many techniques require prior information to infer the trajectory, such as a starting or root cell [8, 9]. Prior biological information can help the method to find the correct trajectory but on the other hand, incorrect prior knowledge can bias the trajectory. Traditionally, previously known stemness markers are used to identify the starting cell, but it is not always feasible due to the high drop-out rate of the scRNA-seq technique. Moreover, stemness markers depend on the tissue and the developmental stage and are not always available for all cases.

In this sense, in order to quantify pluripotency a systems approach methodology was proposed by members of the Teschendorf Lab. They calculated a parameter called network entropy using a protein-protein interaction (PPI) network [10]. In following works the group deepened the research on quantifying stemness, proposing different alternatives to compute entropy [11, 12, 1]. Authors state that differentiated cells have certain specific pathways activated leading to low entropy levels whilst pluripotent cells display a broad pattern of signalling pathways activated and do not express any preference for any particular lineage. In terms of single cell transcriptomics data, this translates as more uniform gene expression profiles, resulting in high entropy levels.

Other publications address entropy without utilizing a PPI network as a scaffold. For instance, StemID aims to identify stem cells by using a score that combines the cluster median entropy and the number of inter-cluster links that define the topology of the lineage tree [13]. Another variation is SLICE, a Shannon entropy-based algorithm with some implementation modifications [14].

On the other hand, there are few techniques to quantify pluripotency that are not based on entropy. For example, Palmer *et al*. derived a stemness gene expression signature and used it to compute a stemness index over gene expression microarray samples [15]. They utilized the projection of the coordinates of an expression profile onto the first principal component of the gene space defined by the stemness gene signature as a relative measure of stemness.

In 2020 Gulati *et al*. developed a computational framework called CytoTRACE to identify stem cells using scRNA-seq data [2]. They found that the gene counts are generally correlated with the state of differentiation. As scRNA-seq was designed to capture gene expression, they suggested determining the genes that correlate with the gene counts and creating a dataset-specific gene count signature (GCS). The authors evidenced a limitation in the case of quiescent stem cells. CytoTRACE is not useful for identifying quiescent stem cells due to their reduced metabolic activity and low RNA content, as is the case for hematopoietic stem cells.

Lately, increasingly information about protein interactions and their role in biological processes is available. In this work we exploit the fact that interactions between proteins underlie cell phenotypes [16]. With the widespread use of high-throughput sequencing technologies, many methodologies to integrate this type of data and network-based biological strategies have been implemented [11]. Here, we use the information of a PPI network to identify stem and progenitor cells. In this way, we define a score, which we call differentiation activity, that quantifies how active the differentiation PPI network is in each cell based on its expression profile and the set reactions involved on this PPI. This tool was implemented as an R package named *ORIGINS*.

### ORIGINS: activity computation

Gene Ontology (GO) provides structured, controlled vocabularies and classifications associated with molecular functions, biological processes and cellular compartment [17]. Exploring GO annotations underlying a set of Differentially Expressed Genes (DEG) for insights into potential experimental meanings has become a widespread practice. Alternatively, to identify differential biological processes (BP) across a cell population we propose a strategy that exploits the set of biochemical reactions involved in the biological function of interest rather than several DEGs. In this sense, we build a protein-protein interaction (PPI) network associated with the gene products involved in Cell Differentiation BP (BP-GO:0030154) as putative biochemical reactions.

To this end we consider 11,582 proteins from H. sapiens associated to BP-GO:0030154 listed at QuickGo database [18], which include 87 child BP terms, these proteins constitute the nodes of the network. Further, we also consider the biochemical interactions listed in Pathways Commons (version 12), that integrate 2,424,055 interactions from 22 databases [19]. From this set of interactions we select the 191,072 interactions that involve only human proteins associated with BP-GO:0030154, we do not take into account interactions involving chemical compounds nor other not-protein molecules. These PPI constitute the edges of the network and can be represented by an adjacency matrix *A*.

After building the PPI network associated with Cell Differentiation BP we define how to compute its activity level from a given expression profile. We consider that the activity level of the pathway is the accumulation of the biochemical reactions occurring in the pathway. Based on the Law of mass action for elementary reactions, we estimate the probability that a reaction will occur as the product of reactant concentrations, without considering the stoichiometric details. This approximation allows us to estimate the contribution of a PPI network edge between nodes *a* and *b*, *a* ↔ *b*, solely from the expression profile as: *x_a_* × *x_b_*, where *x_a_* and *x_b_* are the expression levels associated with proteins A and B, respectively. Therefore, for a given expression profile {*x_i_*} a weighted edge matrix is defined *W_ij_* = *A_ij_x_i_x_j_*, where *i* and *j* = 1, 2, …, *N_g_* and *N_g_* is the number of genes in the pathway. Thus, the activity level associated to Cell Differentiation BP can be defined as:

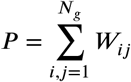

For sc-RNAseq data each cell *k* has an expression profile associated and the corresponding weighted edge matrix *W^k^* from which the activity level *P^k^* associated to Cell Differentiation BP of the *k*th cell can be computed. Finally, activity levels are scaled so that activity takes values between 0 and 1 as follows: 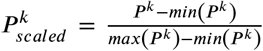, where *min* (*P^k^*) and *max* (*P^k^*) are the minimum and maximum activity levels among all cells.

### Validation

We tested the performance of our proposed methodology to quantify pluripotency using four human data sets: breast epithelium [20], colon epithelium [21], bone marrow (hematopoietic cells) [22] and lungs [23]. Cell types were already annotated and provided as metadata for all datasets except the breast sample, which was annotated by ourselves according to the original publication[20]. For more details on breast cell annotation go to section *Application to human mammary epithelium*. Below is a brief description of the data sets.

#### Breast epithelium

Data from human breast epithelial cells that is publicly available in the GEO database (GSE113197) [20] was used. We utilized a healthy adult sample (Ind4).

#### Colon epithelium

Data was downloaded from the GEO database (GSE125970). We used an adult human colon sample (Colon-2) to benchmark our algorithm.

#### Hematopoietic

Publicly available scRNA-seq data of hematopoietic progenitors from human bone marrow was used. Raw data is available in the GEO database under the accession code GSE117498 [22]. We performed the pluripotency quantification using data from Donor A.

#### Lungs

The scRNA-seq data included 19 lung samples, we focused on one of the five adult donors (D122), a 32 years old healthy male. Data was downloaded from the [cellxgene Data Portal](https://cellxgene.cziscience.com/collections/625f6bf4-2f33-4942-962e-35243d284837). Raw data is also available in the GEO database (GSE161383) [23].

We compared the performance of ORIGINS with two previously existing methodologies specifically developed for scRNA-seq data: the Signaling Entropy Rate (SR) from LandSCENT [1] and CytoTRACE [2]. Both algorithms are publicly available as R packages and were downloaded and installed from their official repositories. In addition, we propose a quick approximation of differentiation activity. We estimated the 2000 top highly variable features (HVF) of the expression matrices using the Seurat function FindVariableFeatures() with selection.method = vst. Thus, we obtained a reduced matrix for each sample and we computed the activity on the normalized reduced matrix. This quantity is referred as activity HVF (ORIGINS).

All the code was implemented in R version 4.1.2 and the main packages used were LandSCENT version 0.99.5, CytoTRACE version 0.3.3 and Seurat version 4.1.0. The computer specifications were Kernel Version 5.13.0-30-generic, processor 12 × Intel® Core™ i7-8700 CPU @ 3.20GHz and 16 GiB of RAM.

The different pluripotency scores calculated for the breast sample are presented in the UMAP space in Figs.1A-D. We investigated how these parameters vary according to cell types. Basal cells presented the highest average levels of SR (LandSCENT), activity (ORIGINS) and its approximation (Figs.1E, G and H). This was expected since in the article where data was published the authors suggested the presence of breast stem cells within the basal population [20]. In the work where LandSCENT is presented, the breast sample was also used and the authors found that the majority of the multipotent cells are basal [1]. In the same way, it would be expected that L1 is in second place because they are more immature cells than L2, however this was only evidenced for activity (ORIGINS) and its approximation. The CytoTRACE score did not coincide with this order, it was found that on average the L1 cluster has the highest values followed by L2 and basal (Figs.1B and F).

**Figure 1:**
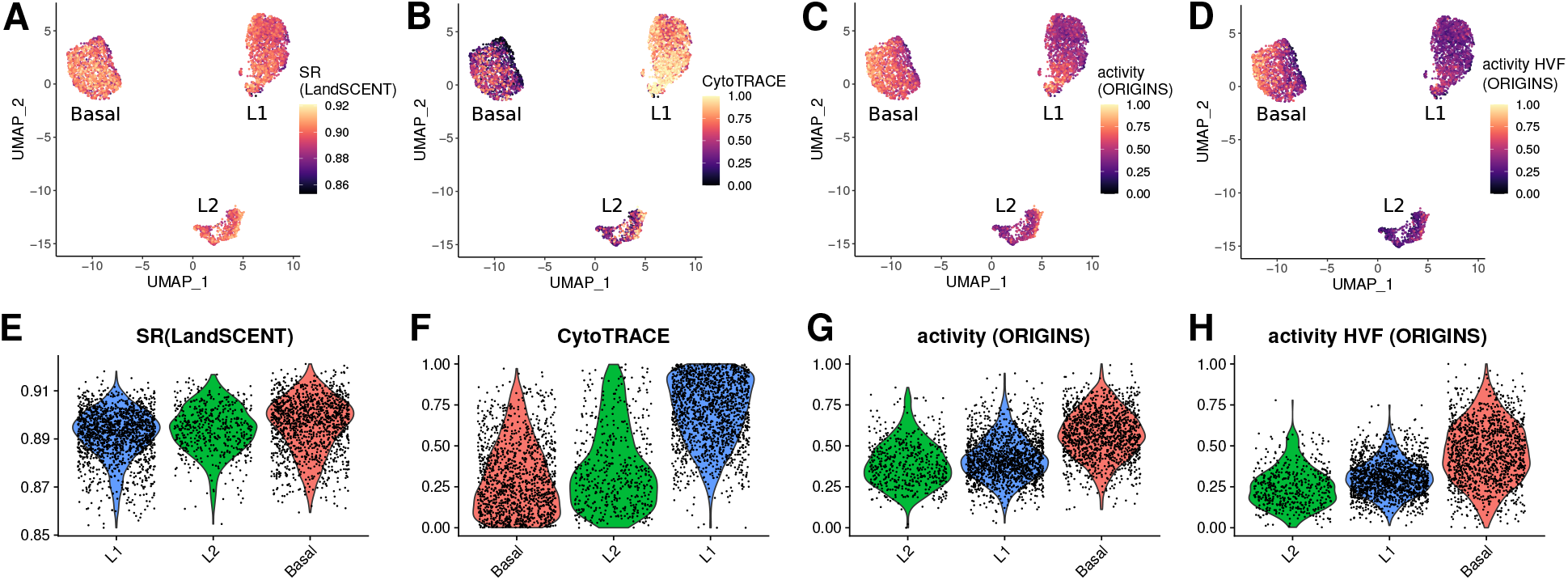
**A-D**: UMAP representation of the breast sample colored by the scores calculated by LandSCENT, CytoTRACE, ORIGINS using all genes and the top highly variable features (HVF). **E-H**: Violin plots of the pluripotency scores per cell type sorted according to increasing values of the mean scores.

In the case of the colon sample, where cells were already annotated (Fig. S1A), the four stemness scores are exhibited in Figs. S1B-E in the UMAP space. All the methods were limited in finding the stem cells, since the transit amplifying cells (TA) showed the highest score values on average and not the stem cells (Figs. S1F-I). In the intestine, stem cells divide asymmetrically, giving rise to another stem cell and a daughter cell called transit amplifying progenitor cell. Transit-amplifying cells are highly proliferative, they undergo a limited number of cell divisions and eventually differentiate into absorptive (enterocytes) or secretory (mucosal, enteroendocrine, Paneth cell) lineages [24, 25, 26].

The analyzed bone marrow cells were classified as hematopoietic stem cells (HSC), multipotent progenitors (MPP), multilymphoid progenitors (MLP), pre-B lymphocytes / Natural Killer cells (PREB/NK), megakaryocyte-erythroid progenitors (MEP), common myeloid progenitors (CMP) and granulocyte-monocyte progenitors (GMP) [22] as shown in Fig. S2A. The classical hematopoietic model states that hematopoietic stem cells (HSC) give rise to all blood-cells types [27, 28]. HSC cells are predominantly in a quiescent state [29] and can be activated as a response to the organism demand [30]. Self-renewing HSC occupy the apex of the hierarchy and originate different progenitors. Figs. S2B-E depict the calculated scores in the UMAP representation for several progenitor cell types. It should be noted that all the methods agreed that the highest average potency corresponded to the GMP as seen in Figs. S2F-I. HSCs would have been expected to be the most pluripotent, followed by MPPs, and then MLPs and CMPs. However, none of the methods seem adequate to order cells according to pluripotency based on the hematopoietic model, this could be a consequence of the different degree of quiescence and cell commitment of the hematopoietic stem/progenitor cells [2, 31, 32].

The lung is a complex organ that includes several distinct cell types, as seen in the lung sample we used (Fig. S3A). Regarding the lungs hierarchical organization, alveolar type II (AT2) cells are the best described stem cells and give rise to alveolar type I (AT1) cells [33]. In addition, club cells are stem cells that differentiate into ciliated cells [34, 35]. Furthermore, basal [36, 37, 38], club-like and pulmonary neuroendocrine cells were identified as progenitor cells [39]. Figs. S3B-E exhibit the pluripotency scores in the UMAP space calculated for the lung sample. Among the approximately 30 cell types present in the dataset analyzed, AT2/club-like had the highest average SR (LandSCENT) and activity (ORIGINS) scores. The highest activity (ORIGINS) values were observed for AT2 cells, although not the highest mean activity score. All cells of interest, including basal and club cells, ranked in the top 8 mean scores for SR (LandSCENT), CytoTRACE, and activity (ORIGINS) as shown in Figs. S3 F,G and H respectively. The activity approximation, HVF activity (ORIGINS), failed to classify cell types based on the expected pluripotency. This may be because the use of the top 2000 HVF is not enough to provide a good activity approximation, probably due to the great diversity of cells analyzed (Fig. S3I).

#### Correlation with other methods

The Pearson correlation coefficient between the methodologies was computed for all data sets as shown in Fig.2. All quantities were positively correlated. Taking into account the four data sets analysed, the average correlation coefficient between activity (ORIGINS) and SR (LandSCENT) was around 0.77, between activity (ORIGINS) and CytoTRACE 0.44, between SR (LandSCENT) and CytoTRACE 0.63 and between activity (ORIGINS) and its approximation activity HVF (ORIGINS) 0.67.

**Figure 2:**
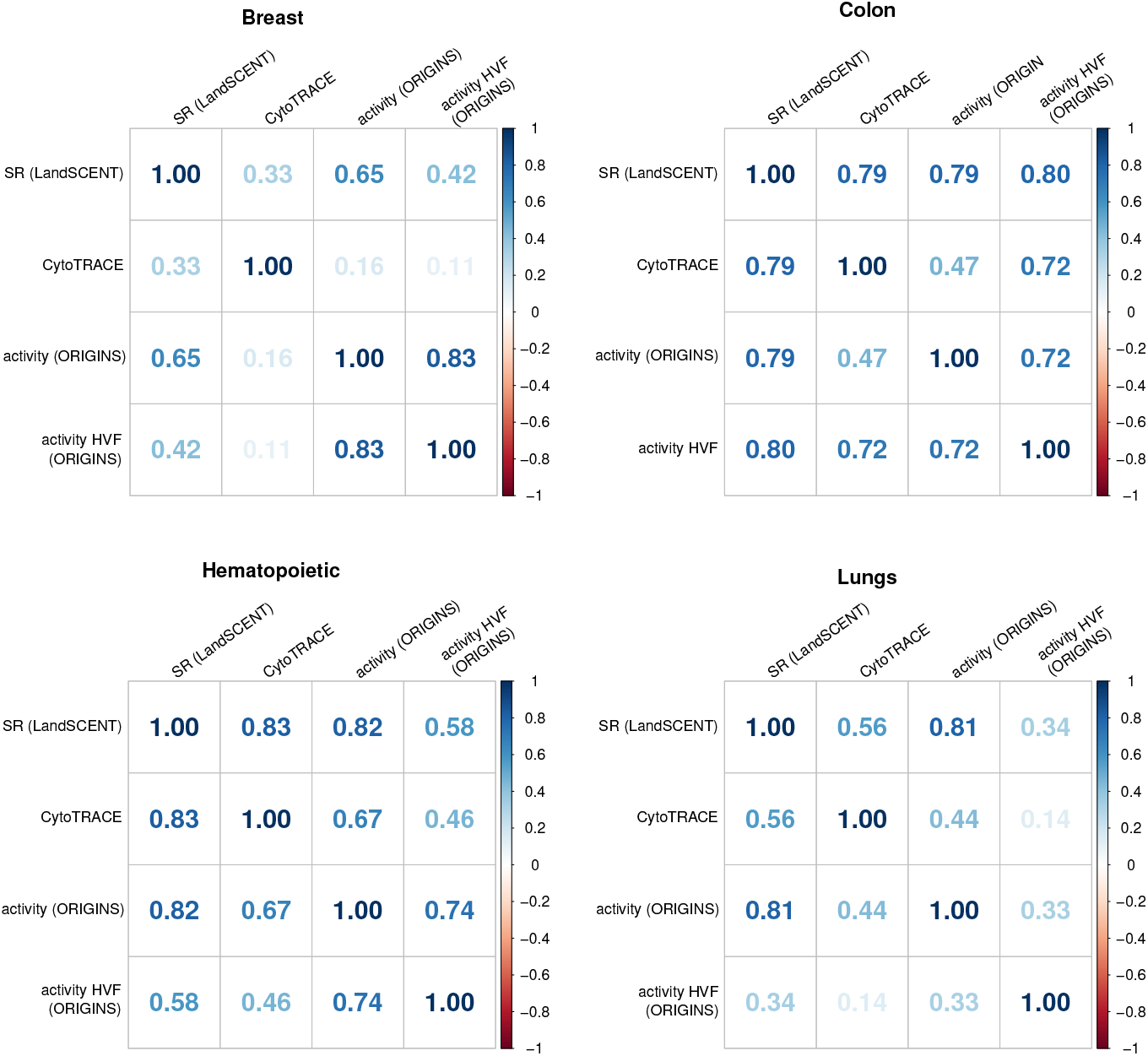
Correlation matrices between all applied methodologies, SR (LandSCENT), CytoTRACE, activity (ORIGINS) and activity HVF (ORIGINS) for all data sets.

#### Efficiency

The elapsed real time for all algorithms and samples are reported in Table 1. On average, SR (LandSCENT) took approximately 25% longer than activity (ORIGINS) and CytoTRACE took less than 1%. As expected, activity HVF (ORIGINS) took less than 2% than activity (ORIGINS).

**Table 1.** Computational time comparisson between SR (LandSCENT), CytoTRACE, activity (ORIGINS) and activity HVF (ORIGINS).

|  | SR (LandSCENT) | CytoTRACE | activity (ORIGINS) | activity HVF (ORIGINS) |
| --- | --- | --- | --- | --- |
| Breast epithelium | 3.43 hrs | 28.27 secs | 2.95 hrs | 3.00 mins |
| Colon | 4.11 hrs | 44.19 secs | 3.11 hrs | 3.11 mins |
| Hematopoietic | 8.12 hrs | 59.95 secs | 6.24 hrs | 4.09 mins |
| Lungs | 5.37 hrs | 35.00 secs | 4.71 hrs | 3.19 mins |

#### RAM usage

LandSCENT was the most memory demanding program. For example, an extra 6.4 Gb of RAM was required for the breast data set, while CytoTRACE needed an additional 5.8 Gb. This makes it difficult to quantify pluripotency for typical size samples (a few thousand cells) using standard personal computers. In this aspect, ORIGINS significantly outperforms the other programs as it does not require additional RAM other than the vector where activity is stored, for the same data set the memory allocated by this vector was 26.6 Kb.

#### Simplicity and biological foundation

The main concept underlying the activity of the differentiation PPI network is relatively simple. Briefly, the differentiation activity of a cell is proportional to the sum of all the weights of the differentiation PPI network. The weights (edges) associated with two transcripts (nodes) are approximated as the multiplication of the expression levels of the associated proteins according to the law of mass action. Thus, an edge weight is greater if both nodes are highly expressed and vice versa. By adding all the edges of this differentiation network, we can quantify how active this network is.

#### ORIGINS can handle zero-containing expression matrices

Normalized expression matrices often have null elements (zeros), such as the obtained by using the R Seurat package normalization. Unlike the Signaling Entropy Rate (SR) [1], ORIGINS has no problem with the presence of zeros in the expression matrix. The user can provide any normalized non-negative expression matrix.

#### User friendly

The algorithm is easy to use. By typing four lines of code in R, the differentiation activity can be calculated:

~~~
**install** . **packages** (“remotes”) *if remotes package not installed*
remotes : : *install*_github (“danielasenraoka/ORIGINS”)
**library** (ORIGINS)
**diff**_activity <-activity (**expression_matrix**, differentiation_edges)
~~~

### Application to human mammary epithelium

We applied our methodology, ORIGINS, to identify stem cells in the human mammary gland. Below we describe all the steps performed, from the raw data to the inference of the differentiation trajectory. We used a scRNA-seq data set of human breast epithelial cells that is publicly available in the GEO database (GSE113197) [20]. This data set was acquired using the 10× Genomics Chromium platform. It included approximately 25000 cells from four nulliparous women aged between 17 and 36 years denoted as Individuals 4 to 7 (Ind4-7). In a previous work the Individual 4 sample (Ind4) was used to quantify pluripotency using LandSCENT [1], so for comparative purposes we will describe our analysis in detail for this donor.

#### Data analysis workflow

We performed the Seurat pipeline for scRNA-seq data analysis. We downloaded the UMI count matrix, cells and features were filtered to reduce noise and eliminate redundancies. Data was normalized and dimensionality reduction was performed (Figs. 3A and B). Clustering and differential expression analysis allowed us to annotate the cells (Figs. S4A and B).We identified three main clusters and concluded that these corresponded to a basal myoepithelial cell type, a luminal immature secretory and immune-related cell type, and a luminal mature hormone-responsive cell type. The identified cell types are in accordance with the original work where the data was published [20] and other subsequent works [1]. In line with the original notation we will refer to them as Basal, Luminal1 (L1) and Luminal2 (L2). The workflow used is described in detail below.

**Figure 3:**
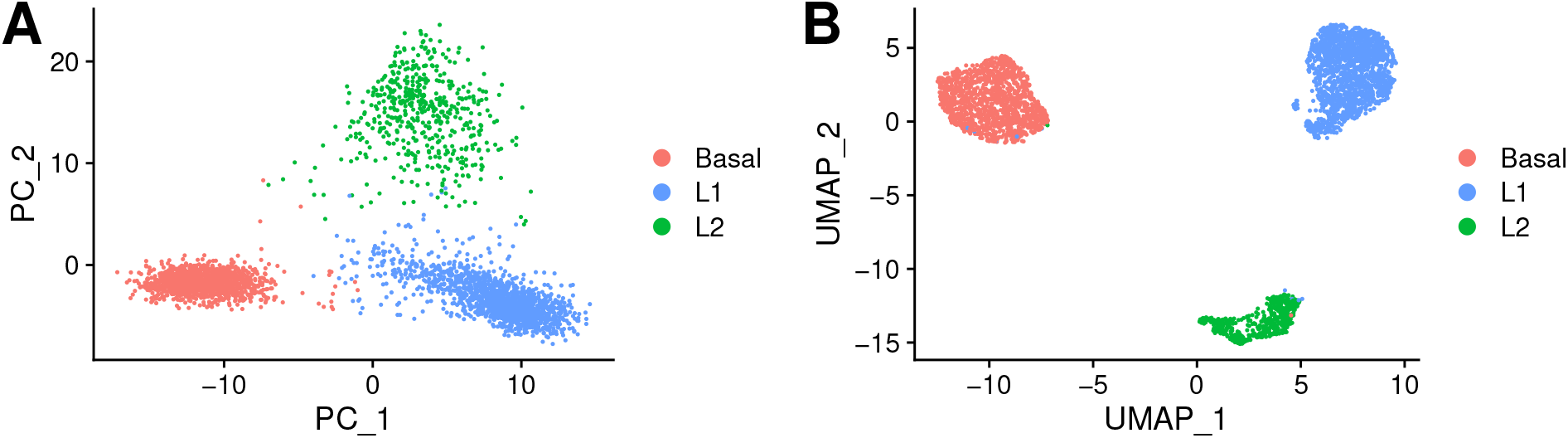
Breast sc-RNAseq data in the PCA (**A**) and UMAP (**B**) space colored by cell type.

##### Filtering

We trimmed cells that expressed atypical amounts of features, either too many or too few, to eliminate doublets or multiplets and debris or empty droplets. We also filtered out features expressed in few cells, the cutoff value of 3 cells per feature was used. Besides, we removed cells that had a high mitochondrial content to eliminate potentially low quality cells (*>* 5%). Additionally, we filtered out three small non-epithelial clusters (stromal, endothelial and outliers) as was done in the work where data was published [20].

##### Normalization

In order to compare cells among them, it is necessary to carry out a normalization step. Gene counts are divided by the total counts for each individual cell, scaled (multiplication by a scale factor) and natural-log transformed. We used the *NormalizeData()* function from the R package Seurat. We later performed a step to identify features that exhibit high cell-to-cell variation.

##### Dimensionality reduction

We scaled the data prior to reducing the dimensionality of the data set, so that the mean expression is 0 and the variance is 1 across cells. We then performed PCA and UMAP and visualized the data (Figs. 3 A and B). Three main big clusters were revealed.

##### Clustering

We performed clustering using the Seurat functions FindNeighbors() and FindClusters(). In short, it includes building a KNN graph using the euclidean distance in the PCA space and applying the Louvain algorithm that optimizes the standard modularity function.

##### Differential expression

We carried out differential expression analysis and obtained the gene signatures of all the clusters. We executed non-parametric Wilcoxon rank sum test and a ROC test that returns the power of a classifier. In Fig. S4A we display some markers in the UMAP space and in Fig. S4B a heatmap is shown to portray the differential expression among clusters.

##### Cell annotation

The previous step provided a gene signature for each cell cluster. Three epithelial cell types were identified. The orange/pink cluster in Figs. 3A and B differentially expressed keratin coding genes such as KRT14 (Fig. S4A KRT14), KRT5 and KRT17. This cluster also displayed smooth muscle related genes, e.g., ACTA2 and MYLK and was labeled as a myoepithelial basal cluster. The two remaining cell clusters were were both positive to KRT18 gene expression and identified as luminal cell types. The blue cluster in Figs. 3A and B exhibited high expression levels of SLPI and LTF (Figs. S4A SLPI and LTF), which are the typical Luminal Progenitor markers. Thus, we annotated this cluster as Luminal 1 (L1). The green cluster in Figs. 3A and B could be identified as a luminal mature cell type because of the differentially expressed ANKRD30A gene (Fig. S4A ANKRD30A). Another gene marker highly expressed was AREG, a central factor in estrogen action and ductal development of the mammary glands. AGR2, which is a hormone responsive gene, was over-expressed too. Overall, this cluster was labeled as Luminal 2 (L2) and is associated with a hormone responsive function.

#### Pluripotency quantification using ORIGINS

Differentiation PPI network activity was computed over the gene expression matrix. The normalized expression matrix used to calculate the activity was not the one provided by Seurat because this procedure returns a matrix that has null elements. Since our goal is to compare our algorithm performance with other methods, and LandSCENT does not accept expression matrices with null elements as input, we applied a different normalization. We followed the normalization on the LandSCENT tutorial which sets an offset value of 1.1 before log-transformation to avoid having zero values.

Differentiation activity can be visualized in the PCA and UMAP spaces in Figs. 4A and B. The highest levels of activity were found within the basal group. This is in agreement with the results of the work in which the data was originally published, as the authors found a group of basal cells with stemness capacity [20]. Similarly, in the LandSCENT publication, the authors found a higher percentage of multipotent cells within the basal cluster, although they also observed high levels of SR in luminal cells in close proximity to the basal cluster [1].

**Figure 4:**
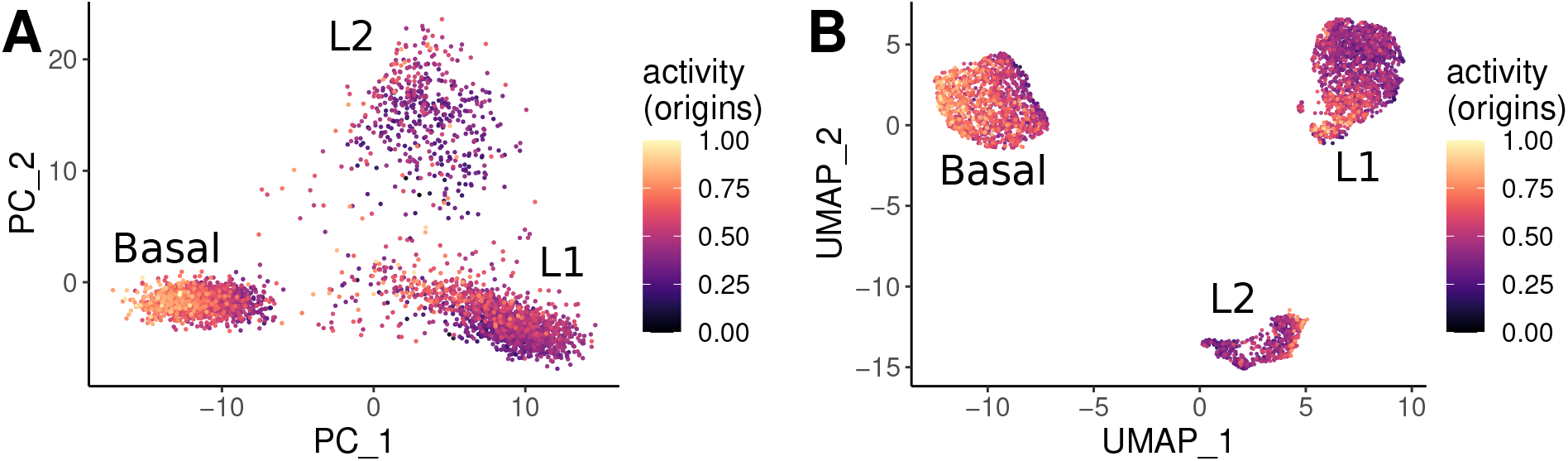
Breast sc-RNAseq data representation in the PCA (**A**) and UMAP (**B**) spaces colored by the differentiation PPI network activity (ORIGINS).

#### Trajectory inference

Trajectory inference are single cell transcriptomics techniques that infer lineages structures. Essentially, these computational methods order cells based on their expression similarities along a “temporal” variable called pseudotime. In agreement with Saelens *et al*. [4] we consider that, among the large number of tools available, Slingshot [40] is one of the simplest, most robust and best documented. For this reason we run Slingshot, an R package, on the breast data set. Slingshot works in two steps. First, it infers the global lineage structure using a cluster-based minimum spanning tree. In a second instance, it infers pseudotime for each lineage using simultaneous principal curves. Like many methods, Slingshot requires the prior definition of the trajectory origin, namely, the stem or progenitor cells.

We set the root as the cells with the highest differentiation PPI network activity, located within the basal cluster. We discovered that the trajectory progresses from the basal starting point and bifurcates into the L1 and L2 cell types, going through an intermediate state (Figs. 5A and B). Furthermore, this group of luminal precursor cells located within the L1 cluster has moderately high activity levels (Fig.4A). At this point the trajectory branches and heads towards the terminal cells L1 or L2 into two separate lineages. The results obtained by performing trajectory inference supports previous works where stem/progenitor breast cells where suggested as bi-potent cells that can originate differentiated basal and luminal cell types [20, 1]. We refer to the lineage starting at the basal cluster and ending at cluster L1 as Lineage 1 and the one ending at L2 as Lineage 2.

**Figure 5:**
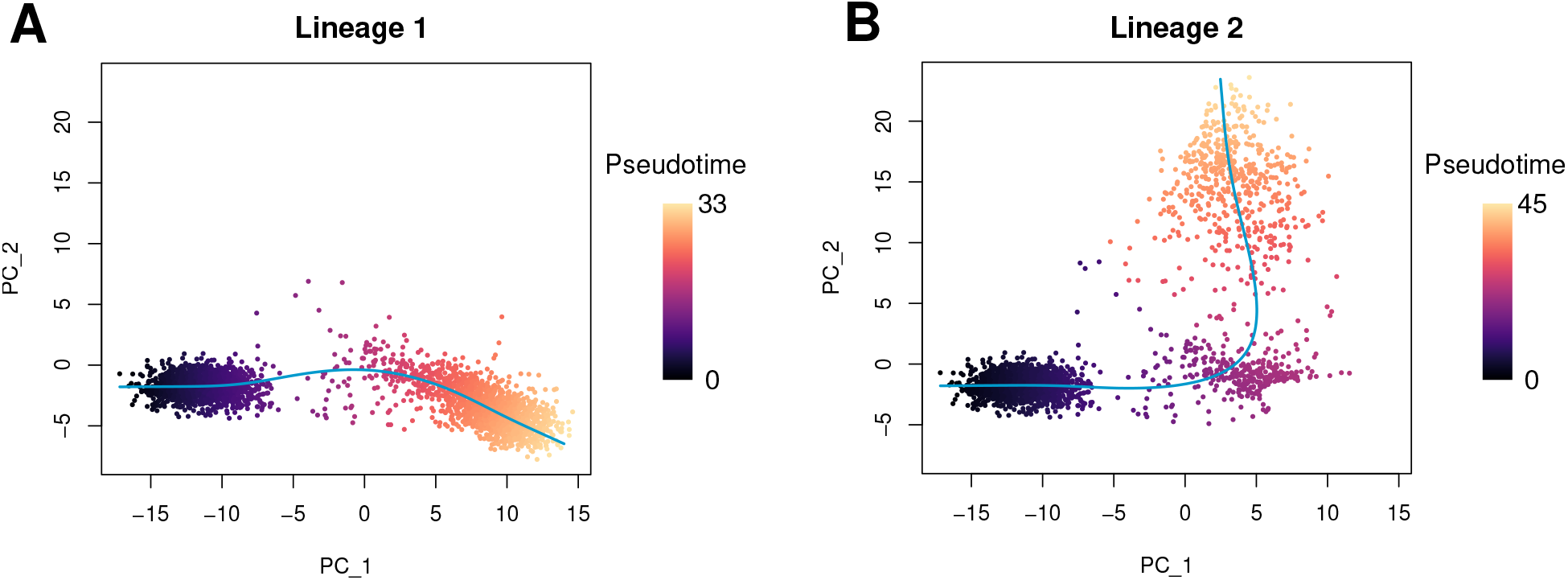
Breast sc-RNAseq data representation in the PCA space colored by pseudotime. Performing trajectory inference revealed a bifurcating pathway. **A**: The first branch, lineage 1, leads to the L1 cell type. **B**: The second branch, lineage 2, ends at the L2 cell cluster.

We next performed differential expression analysis along the inferred trajectory to both lineages. Thereby, we obtained the genes whose expression changed the most along the trajectory, which can be visualized by the heatmaps in Figs. S5A and B. For that, we used the R Bioconductor package tradeSeq [41]. Briefly, tradeSeq fits a negative binomial generalized additive model (GAM) to model the relation between gene expression and pseudotime and then tests for significant relationships between gene expression and pseudotime.

In summary, we show a complete workflow to perform trajectory inference with sc-RNAseq data. We first identified the root cells using our algorithm (ORIGINS) and then used SLINGSHOT to unveil the differentiation process of the breast epithelium. SLINGSHOT provided a bifurcated trajectory, supporting the theory that breast stem cells are bi-potent. Furthermore, our analysis allowed us to identify the genes that drive the differentiation process.

## Data availability

Data analyzed in this work is publicly available on the [GEO database](https://www.ncbi.nlm.nih.gov/geo/) under the accession codes: GSE113197 for breast epithelial cells, GSE125970 for colon epithelial cells, GSE117498 for hematopoietic bone marrow cells and GSE161383 for lung cells.

## Code availability

ORIGINS is freely available as an open source R package from the GitHub repository: https://github.com/danielasenraoka/ORIGINS.

## Acknowledgements

This research was supported by CONICET, Argentina (PIP 1748). NG and LD are researchers and DS is a PhD fellow at CONICET, Argentina.

## Author contributions

LD conceptualized the paper. NG, LD and DS developed the the structure and design of the manuscript. LD and DS developed the analysis and algorithms and wrote the manuscript. DS implemented the software. NG, LD and DS revised the manuscript and approved the final version.

## Supplementary Materials

Figs. S1-S5.

