## Supplementary Materials for "ORIGINS: a protein network-based approach to quantify cell pluripotency from scRNA-seq data"

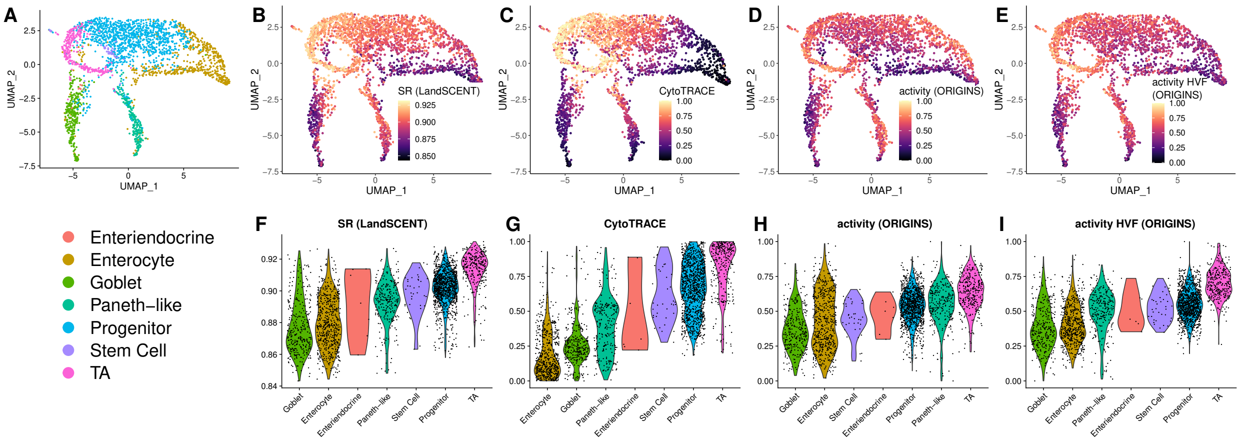


**Figure S1: A-E**: UMAP representation of the colon sample colored by cell types and the scores calculated by LandSCENT, CytoTRACE, ORIGINS using all genes and the top highly variable features (HVF). **F-I**: Violin plots of the pluripotency scores per cell type sorted according to increasing values of the mean scores.

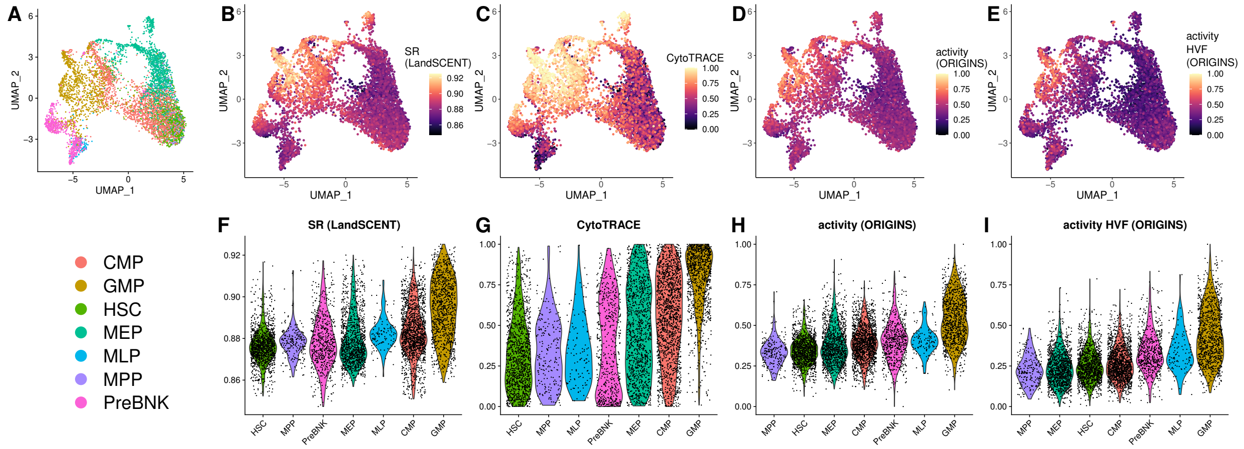


**Figure S2: A-E**: UMAP representation of the bone marrow sample colored by cell types and the scores calculated by LandSCENT, CytoTRACE, ORIGINS using all genes and the top highly variable features (HVF). Cell types are abbreviated as follows: hematopoietic stem cells (HSC), multipotent progenitors (MPP), multilymphoid progenitors (MLP), pre-B lymphocytes / Natural Killer cells (PreBNK), megakaryocyte-erythroid progenitors (MEP), common myeloid progenitors (CMP) and granulocyte-monocyte progenitors (GMP). **F-I**: Violin plots of the pluripotency scores per cell type sorted according to increasing values of the mean scores.

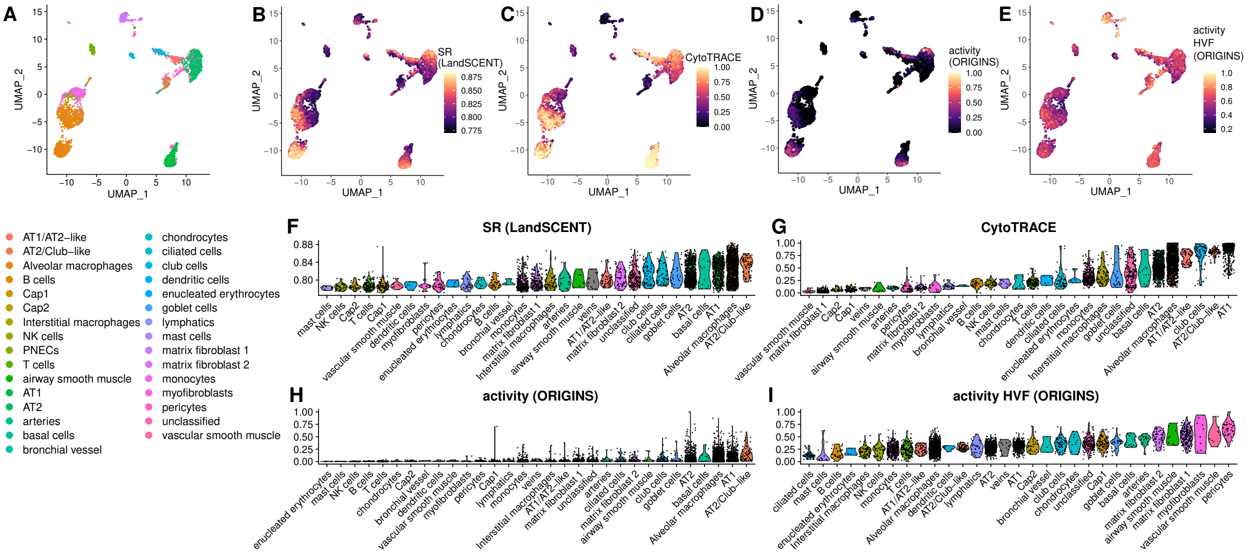


**Figure S3: A-E**: UMAP representation of the lung sample colored by cell types and the scores calculated by LandSCENT, CytoTRACE, ORIGINS using all genes and the top highly variable features (HVF). **F-I**: Violin plots of the pluripotency scores per cell type sorted according to increasing values of the mean scores.

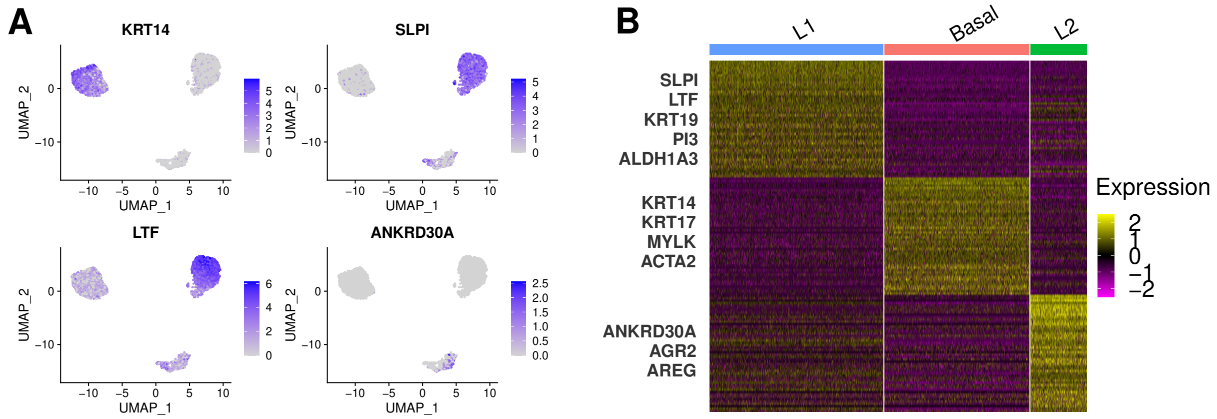


**Figure S4: A**: UMAP representation of the breast sample with typical markers highlighted. **B**: Heatmap displaying some differentially expressed features for the three cell types.

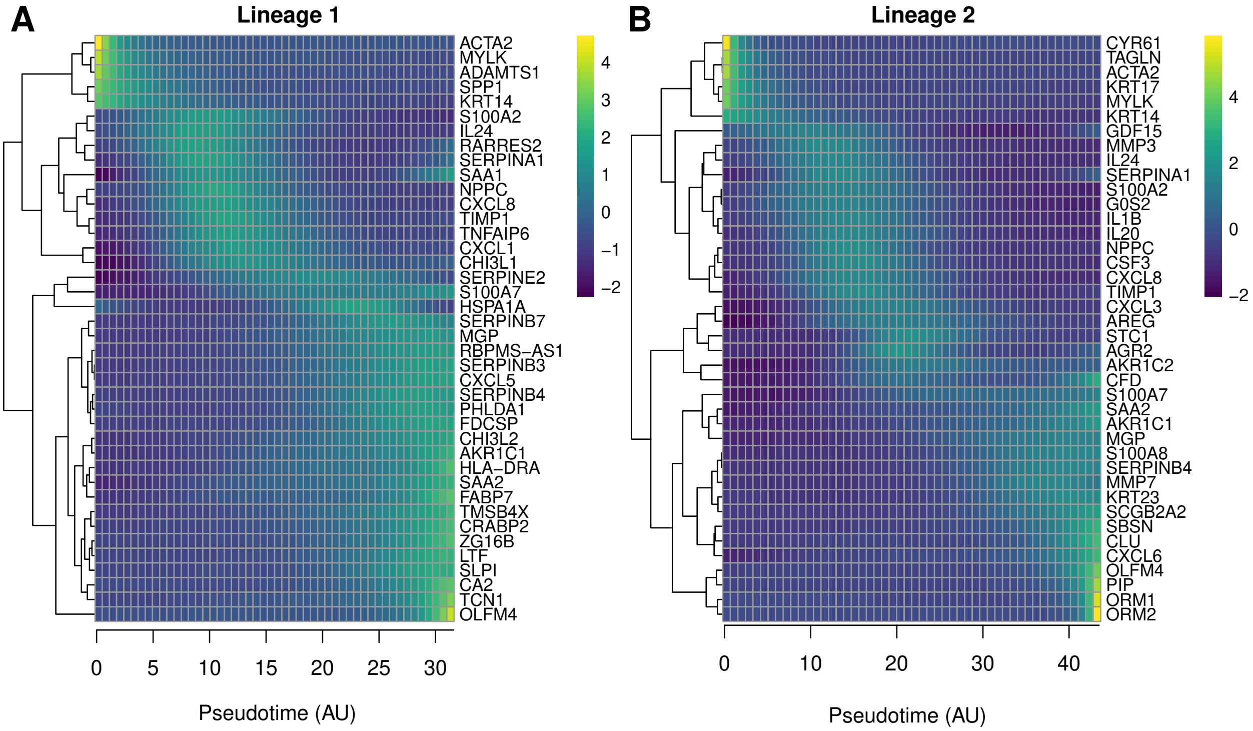


**Figure S5:** Top 40 genes differentially expressed along the trajectory for Lineage 1 (**A**) and Lineage 2 (**B**) of the breast sample.
